# Different categories of fluorescent proteins result in GEVIs with similar characteristics

**DOI:** 10.1101/2020.05.06.081018

**Authors:** Jelena Platisa, Zhou Han, Vincent A. Pieribone

## Abstract

The latest generation of genetically encoded voltage indicators (GEVIs) is significantly advancing our ability to study electrical activity from large numbers of identified neurons. The further refinement of the technology will contribute to our understanding of behavior-evoked information perception, transfer and processing on a cellular level across brain regions. The development of GEVIs relies on synthetic biology which includes rational and random modifications of indicator sequence. One strategy in GEVI design is based on creating chimeras between voltage sensitive protein domains (VSDs) and fluorescent proteins (FPs). However, in this design scenario, the mechanistic details of voltage-induced fluorescence change that would inform rational design and improvements of GEVIs are still largely missing. Here we preformed a systematic study of how nature of the FP and altering the insertion site affects the characteristics of *Ciona intestinalis* voltage-sensitive phosphatase-based GEVIs. Surprisingly, we found that regardless of vast difference in phylogenesis, biochemical properties, fluorophore structure, sequence and excitation/emission spectra between FPs, the resulting GEVIs exhibit virtually identical decrease in fluorescence intensity in response to depolarization. These results stand in strong contrast to studies demonstrating that small numbers of targeted mutations in the FP sequence cause dramatic changes in both signal size and polarity.

## Introduction

Recording electrical activity of neurons provides the most direct readout of functional brain activity. To understand the massively parallel and yet distributed processing attributes of complex nervous systems, it will be necessary to monitor behaviorally associated neuronal electrical activity in large numbers of networked neurons. The advent of genetically encoded, protein-based voltage indicators (GEVIs) has made simultaneous recording of electrical activity in millions of identified neurons plausible for the first time. GEVIs that are showing the most promise for practical use are designed as chimeras combining voltage sensitivity of a phosphatase or microbial opsin with fluorescent protein(s) ^1–3^. The combination of different voltage sensors, different fluorescent proteins, and different construction designs has resulted in numerous GEVIs that differ in sensitivity, speed of response, and optical characteristics (reviewed in ^4, 3^). However, regardless of the increasing number of physiological studies in which GEVIs are used in various model organisms ^5–13^, we are still lacking variants that can be used across all experimental conditions (e.g. type of activity vs. imaging system). For example, recent studies have demonstrated that opsin-based GEVIs (i.e. Ace2N-2AA-mNeon, MacQ-mCitrine) do not work under multiphoton illumination, the technique of choice for deep tissue imaging necessary in highly scattering media such as the mammalian brain ^8, 14, 15^.

Mechanistic studies have shown that the voltage-dependent fluorescence change in opsin GEVIs depends on changes in the protonation state of the Schiff-base chromophore ^16, 17^. Further, in indicators based on opsin-FP chimeras, this voltage induced change in opsin fluorescence is transferred to a fluorescence change in a FRET compatible partner FP ^2^. In contrast, our understanding of the output modulation mechanism(s) for VSD-FP(s) GEVIs (i.e. ArcLight, ASAP, Marina, Butterfly, etc.) are much more vague. The modulation of fluorescence in VSD-based GEVIs has been achieved with a single attached fluorescent protein ^18, 19^, FRET capable pairs of FPs ^1, 20–22^ or circular permuted FPs ^23–25^. There is clear mechanistic rationale and experimental biophysical support for modulation of fluorescence between FRET pairs (i.e. Butterfly) however, the mechanisms(s) by which the fluorescence is modulated in single inserted FP GEVIs (i.e. ArcLight, Marina and ASAP) is unclear. Without atomic structures of GEVIs in different states of transmembrane potential, it will be difficult to propose the mechanisms of fluorescence output change in these indicators. Consequently, studies focused on GEVI operational mechanics are likely to contribute to our ability to use rational manipulations to target promising templates for desirable properties ^19, 26, 27^.

In this work we tested the performance of a series of fluorescent proteins with different spectral characteristic as optical reporters in GEVIs. We examine both the effects of the nature of fluorescent protein and altering the insertion site on the characteristics of *Ciona* voltage-sensitive phosphatase-based GEVIs. Our result show that regardless of wide range of phylogenesis, structural and spectral differences of the FPs, the resulting GEVIs exhibit shared response profile characterized by depolarization-dependent decrease in fluorescence intensity. The presented results are discussed in the light of previous studies where advances in probe performance were made via targeted mutations in fluorescent protein of GEVI ^18, 19^.

## Results

### Fluorescent protein selection

We were interested in testing fluorescent proteins with diverse optical properties and evolutionary background for their potential to be optical reporters of voltage in CiVSD-based GEVIs. We choose FPs derived from several different organisms and with the excitation/emission maximum spanning over visible spectra, ~400nm to ~650nm (Table 1 and Suppl Table 1). For the blue-shifted FP variants, we used jellyfish *Aequorea victoria* GFP (avGFP)-derived fluorescent proteins mCerulean ^28^, eGFP ^29^, ecliptic pHluorin GFP ^30^ and YFP^31^. In addition, we tested recently developed green/yellow fluorescent protein mNeonGreen derived from LanYFP (yellow fluorescent protein) isolated from lancelet *Branchiostoma lanceolatum ^32^.* The fluorescent proteins with the red-shifted emission originated from several different organisms. Marine anemone, *Discosoma striata* derived fluorescent protein dsRED is the parent of the “fruits family” from which we tested mOrange, mOrange2, tdTomato, mTangerine, mStrawberry and mCherry ^33, 34^. The TagRFP-family, FusionRed, and Neptune-family (mNeptune, mNeptune2 and mCardinal) are derivatives of eqFP578 isolated from another anemone, *Entacmaea quadricolor* ^34–38^. mScarlet is highly bright monomeric fluorescent protein that has been developed *de novo* using synthetic approach ^39^. Finally, we tested near-red iRFP713, derived from bacteriophytochrome RpBphP2 isolated from bacterium *Rhodopseudomonas palustris* ^40^. These fluorescent protein were fused to the voltage sensitive domain (VSD) derived from *Ciona intesitinalis* voltage sensitive phosphatase (CiVSP) ^41^. The selection of the insertion site within CiVSD was made based on the previously developed GEVIs: R254 (as seen in GEVI VSFP2.1^1^), S249 (as seen in GEVI Mermaid ^42^) and Q239 (as seen in GEVI ArcLight^18^) (Figure 1).

**Table 1.**
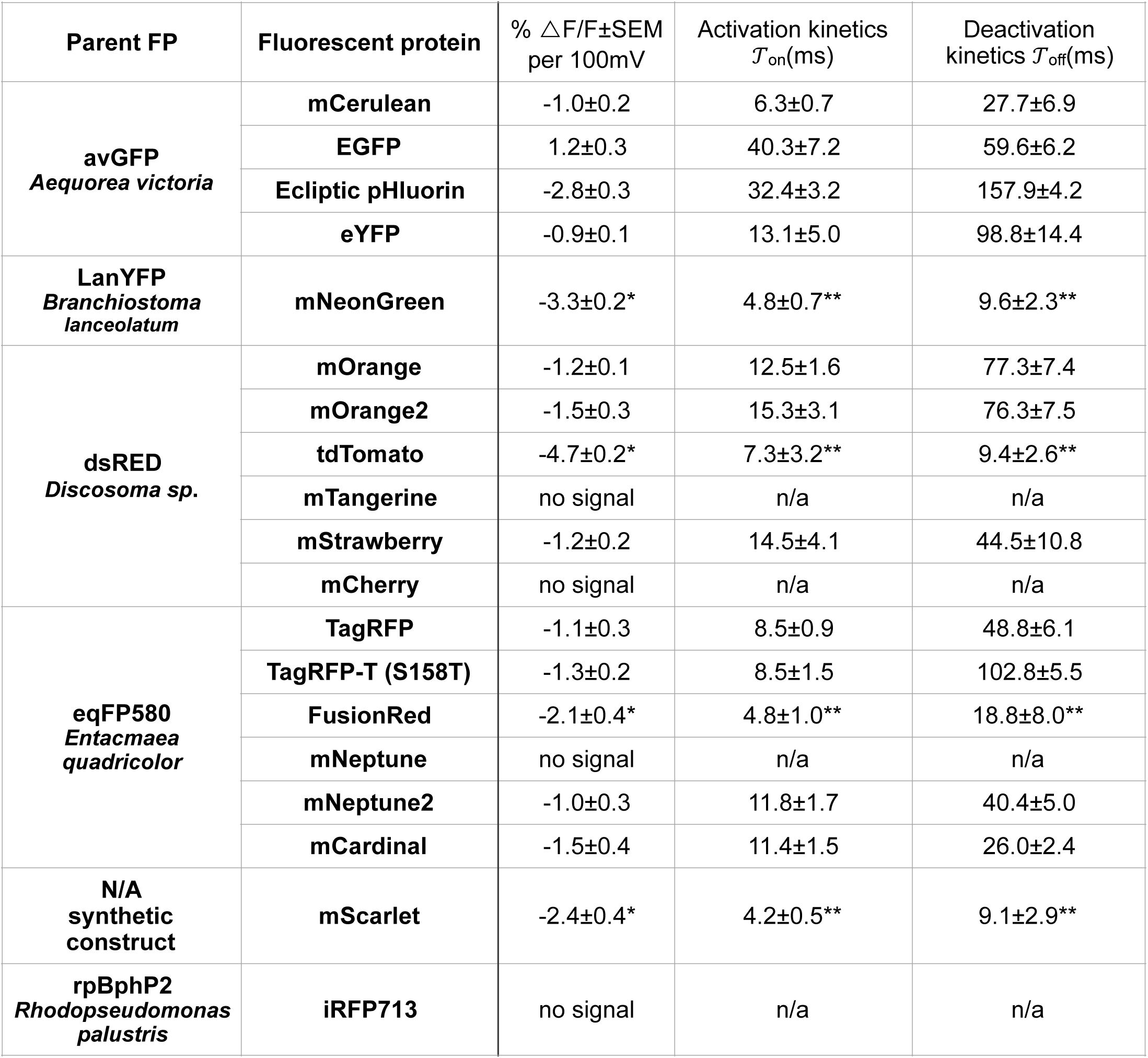
Functional characterization of voltage sensitivity in CiVS-based indicators using various fluorescent proteins. In the majority of constructs, insertion site of FP was residue R249 with linker GDP. The exempts (labeled with * in the table) are constructs with FPs inserted at the residue Q239 tdTomato (linker GDP) and mNeonGreen, mScarlet and FusionRed (linker GNS). For construct characterization, we used concomitant whole-cell patch clamp and optical imaging of transiently expressing HEK293 cells. At least three cells were tested per a construct. The data are shown as mean ± SEM. For majority of constructs, the kinetics of the response was fitted using single exponential curve. Several most sensitive constructs had kinetics that was best fitted using double exponential curve, in which case table values represent the fast component of the activation or deactivation (labeled with ** in the table; mNeonGreen, tdTomato, mScarlet and FusionRed GEVIs).

**Figure 1.**
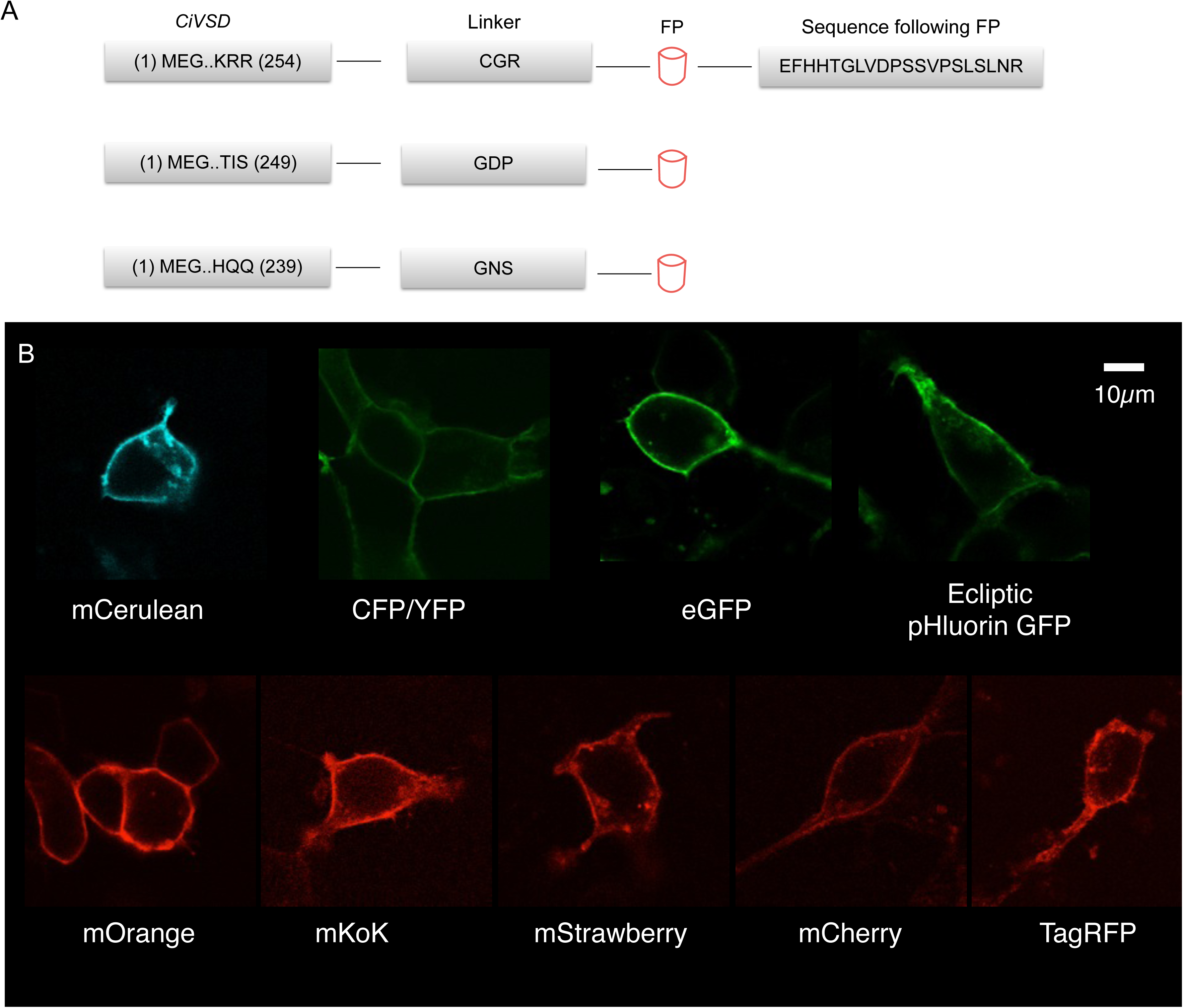
The GEVI design and examples of cellular expression in cultured mammalian cells **A.** The protein sequence of the insertion sites (within voltage sensitive domain of CiVSD) and the linkers for GEVI used in this study The various fluorescent proteins were fused to voltage sensitive domain of *Ciona intestinallis* (CiVSD) at residue R254 with linker CGR, A249 with linker GDP and/or Q239 with linker GDP or GNS. **B.** Cultured mammalian cells (HEK293) transiently expressing GEVIs were imaged using confocal microscopy. All constructs created for this study produced membrane localized bright fluorescence regardless of the fluorescent protein, insertion site or linker composition. The fluorescent proteins are indicated in the panel.

**Figure 2.**
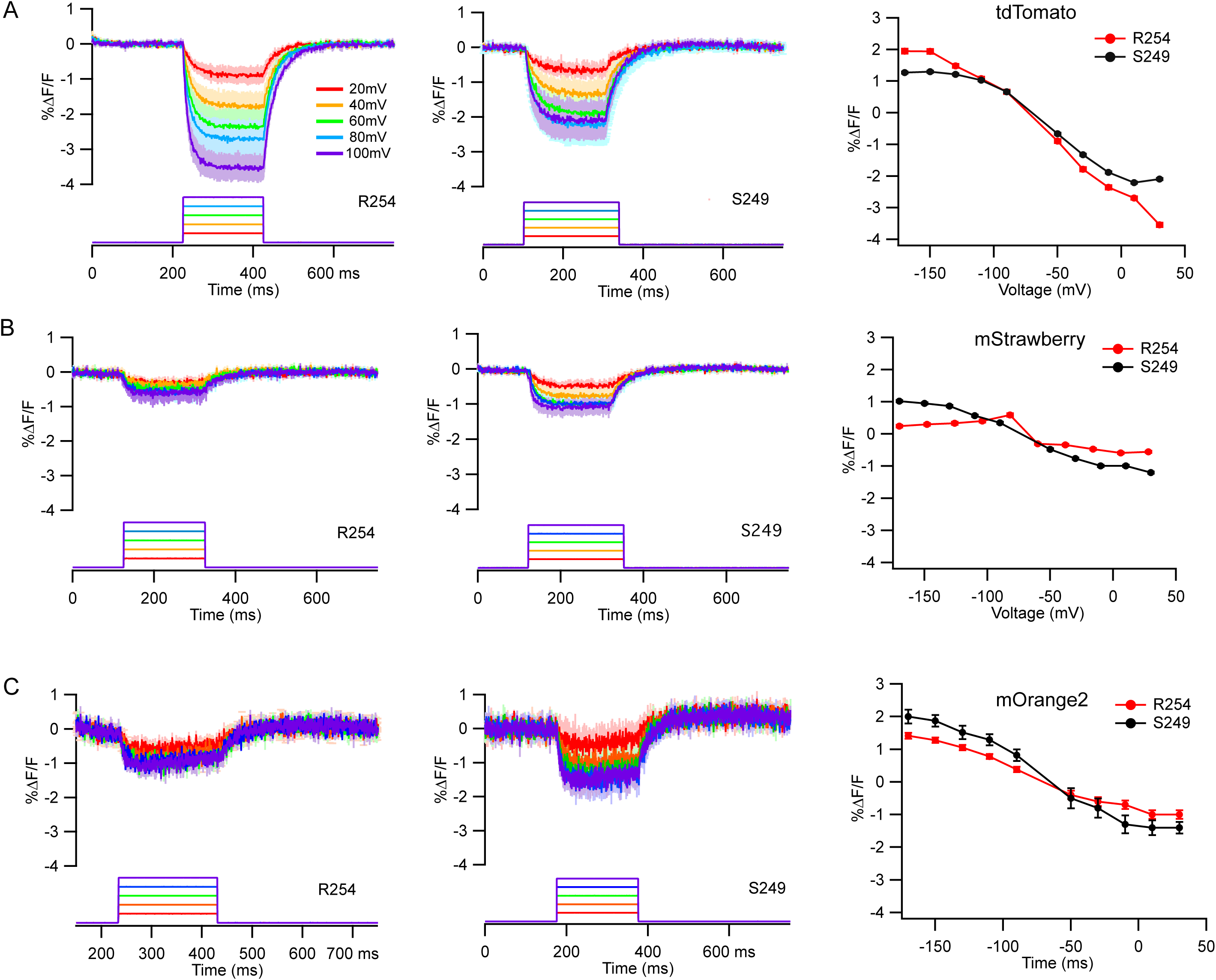
Examples of voltage-dependent fluorescence responses of GEVIs tested in cultured mammalian cells using patch-clamp fluorimetry. All experiments were preformed 24-48h postransfection in HEK293 cells transiently expressing GEVIs. The total of 10 200-250ms long, -/+20mV incremental voltage steps were applied to cells from −70mV resting potential (−170mV to +170 mV). In all panels: left is FP inserted at R254, middle is FP inserted at A249, and right are V/F curves for both. The hyperpolarization steps/responses used for creation of V/F curve are not shown. The signals are unfiltered average of responses from ten trials recorded in three to seven cells. The traces were corrected for the photobleach effect. For color coding of voltage steps/signals see legend in the panel A. The standard error (SEM) traces are shown either in the light color (left and middle panels) or as error bars (right panels). A) tdTomato, B) mStrawberry, and C) mOrange2.

### Functional characterization of potential GEVIs

The potential GEVI constructs were tested using concurrent high speed (1kHz or 2kHz), wide-field imaging and whole-cell patch clamp recordings (100mV depolarization voltage steps) in HEK293 cells transiently expressing potential indicators (24-48 hours post transfection). As shown in Table 1 and 2, the majority of constructs showed relatively slow, depolarization-dependent decrease in fluorescence intensity. The voltage dependent fractional change in fluorescent intensity across tested constructs was ranging from - 0.7 to - 4.7 % ∆F/F for 100mV depolarization steps. Among blue-shifted FPs, the most sensitive indicator was with mNeonGreen fused to VSD at the Q239 residue with fractional change of −3.3±0.2 % ∆F/F for 100mV depolarization step. The only indicator with positive relationship between fluorescence and voltage change was based on eGFP. The eGFP fusion site affected the sensitivity of the indicators, with signal of 0.7±0.1 % ∆F/F at R254 and 1.2±0.3 % ∆F/F at S249 fusion site for 100mV depolarization steps. Among red-shifted FPs, the majority of constructs showed low voltage sensitivity with average fluorescence decrease of ~1% ∆F/F for 100mV depolarization step (Table 1). The most sensitive red-shifted indicator (−4.7±0.2 % ∆F/F for 100mV depolarization step) was with tdTomato fused to CiVSD at the Q239 residue. In contrast, the indicators based on the mTangerine, mCherry, mNeptune and iRFP713 did not produce viable fluorescent signals in response to tested voltage fluctuation. Kinetics of the fluorescence change varied significantly between indicators with different FPs. However, regardless of the FP, there was a noticeable trend that activation rate is faster than deactivation rate (Table 1 and 2). In most cases, the kinetics curves of activation and deactivation were fitted using single exponential curve. Previously reported biexponential kinetics of *CiVSD*-based GEVIs ^43^ was visible in the more sensitive GEVI variants (mNeonGreen, tdTomato, FusionRed and mScarlet).

**Table 2:** Functional characterization of voltage sensitivity of FP GEVIs with different insertion sites. For construct characterization, we used concomitant whole-cell patch clamp and optical imaging of transiently expressing HEK293 cells. At least three cells were tested per a construct. The data are shown as mean ± SEM. For majority of constructs, the kinetics of the response was fitted using single exponential curve. tdTomato constructs were fitted using double exponential curve, and table values represent the fast component of the activation or deactivation (labeled with “**” in the table)

| Parent FP | Fluorescent protein | S249 |  |  | R254 |  |  |
| --- | --- | --- | --- | --- | --- | --- | --- |
| | | % $\Delta F/F \pm \text{SEM}$<br>per 100mV | Activation<br>kinetics<br>$T_{\text{on}}(\text{ms})$ | Deactivation<br>kinetics<br>$T_{\text{off}}(\text{ms})$ | % $\Delta F/F \pm \text{SEM}$<br>per 100mV | Activation<br>kinetics<br>$T_{\text{on}}(\text{ms})$ | Deactivation<br>kinetics<br>$T_{\text{off}}(\text{ms})$ |
| **avGFP**<br>*Aequorea victoria* | **mCerulean** | -1.0±0.2 | 6.3±0.7 | 27.7±6.9 | -1.0±0.1 | 8.0±1.4 | 36.1±6.0 |
| | **EGFP** | 1.2±0.3 | 40.3±7.2 | 59.6±6.2 | 0.7±0.1 | 57.0±11.6 | 132.5±16.2 |
| | **Ecliptic pHluorin** | -2.8±0.3 | 32.4±3.2 | 157.9±4.2 | -0.7±0.1 | 14.0±1.8 | 204.8±57.5 |
| **dsRED**<br>*Discosoma sp.* | **mOrange** | -1.2±0.1 | 12.5±1.6 | 77.3±7.4 | -0.8±0.1 | 5.3±1.3 | 24.0±9.5 |
| | **mOrange2** | -1.5±0.3 | 15.3±3.1 | 76.3±7.5 | -1.1±0.1 | 13.8±4.5 | 114.6±25.6 |
| | **tdTomato** | -2.2±0.5 | 11.1±2.4** | 21.7±7.2** | -3.5±0.3 | 9.5±1.2** | 16.2±3.6** |
| | **mStrawberry** | -1.2±0.2 | 14.5±4.1 | 44.5±10.8 | -0.7±0.2 | 10.6±2.4 | 47.6±5.8 |

Voltage sensitivity of control constructs, FRET-based GEVIs Mermaid and VSFP2.1 tested under the same conditions was comparable to what was reported in original publications (data not shown).

### Insertion site optimization

In previous studies we showed that the insertion site of a FP within the VSD can have profound effects on voltage sensitivity. For example, the sensitivity of the GFP-based indicator ArcLight was doubled by moving the FP closer to the S4-domain of the voltage sensor (~-35% from ~-18%) ^18, 44^. Likewise, initial testing of several blue-shifted (mCerulean, eGFP, and ecliptic pHluorin GFP) and red-shifted (mOrange, mOrange2, tdTomato, and mStrawberry) FPs in this study showed a similar trend with an increase in voltage sensitivity when the FP is positioned closer to the S4-domain of the VSD (S249 vs. R254) (Table 2). Further, we created a series of GEVIs in which the FP tdTomato was fused to the VSP at seven positions (R254, S249, S243, A242, K241, M240 and Q239-Figure 3). Fusing the FP tdTomato closer to the S4-domain resulted in variants that showed an one-fold increase in voltage sensitivity. The highest performing variant, Q239, had a response of −4.7±0.2 %∆F/F compared to −3.4±0.3 %∆F/F (per 100mV depolarization step) seen in the original R254 variant. The effect of insertion site on the sensor kinetics was variable (Table 2 and Figure 3). The fastest variants being with the tdTomato at position S243 (3±1 ms for fast on kinetics; 9±3 ms for fast off kinetics) and Q239 (7±3 ms for fast on kinetics; 9±3 ms for fast off kinetics).

**Figure 3.**
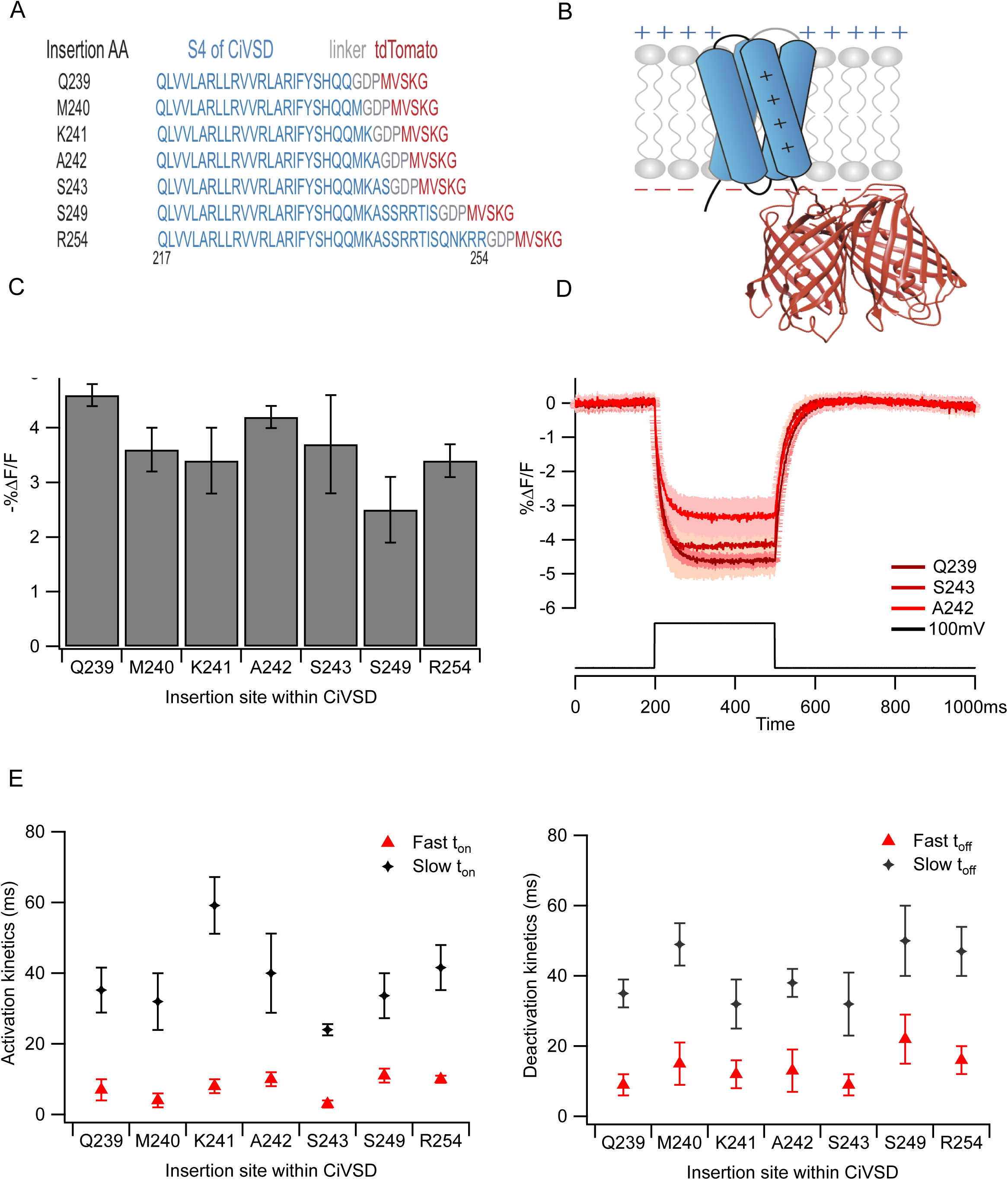
The insertion site modification in the tdTomato GEVI. **A.** Protein sequences of the seven different fusion sites used for insertion of the tdTomato within the voltage sensitive domain of Ciona intestinallis (CiVSD). **B.** Schematic diagram of the tdTomato GEVI design. The transmembrane S1-S4 domains of CiVSD are in blue; the intracellularly localized tandem dimer FP tdTomato is shown in red. **C.** The depolarization-dependent response of seven variants of the tdTomato GEVI. All constructs were tested in transiently expressing HEK293 cells using patch-clam fluorimetry. The 300ms long, +100mV depolarization voltage steps were applied from - 70mV resting potential. Signals are averages of responses recorded in three to seven cells. The error bars are standard error (SEM). **D.** Examples of averaged fluorescence traces recorded from constructs shown in C. For color coding see figure legend. The standard error (SEM) is presented in light color. All traces are averages of ten trials, unfiltered and bleach corrected. **E.** The kinetics of activation (left) and deactivation (right) of fluorescence response. The traces were fitted using double exponential curve.

### Mutagenesis of red-shifted fluorescent proteins

Previously we showed that changes in polarity of outward oriented residues in the *β* barrel (beta sheets 7, 10 and 11) of the pHluorin SuperEcliptic GFP result in increased voltage sensitivity ^18, 19^. In addition, mutagenesis of the residues surrounding the chromophore affected the direction of the voltage response in the GFP-based GEVIs (depolarization dependent increase vs. decrease in fluorescence intensity) ^19^. Recently, this approach was implemented on the red-shifted GEVI Ilom which is based on the dimer FP dTomato ^45^. The introduction of charged residues at the surface of the *β* barrel of dTomato failed to improve the voltage sensitivity. To further examine the potential effect of such manipulations on the red-shifted GEVIs we designed targeted mutagenic libraries of two red-shifted fluorescent proteins, mScarlet and FusionRed. These red FPs were chosen due to their favorable characteristics, mScarlet is synthetically designed monomer with exceedingly high extinction coefficient and quantum yield (100,000 M^-1^ c^-1^ and 0.7), and FusionRed was shown to express superbly in mammalian cells ^37, 39^. In addition, unlike dTomato both FPs are optimized monomers. The native constructs based on mScarlet or FusionRed fused at Q239 to CiVSD showed similar voltage sensitivity of ~-2 %∆F/F per 100mV depolarization step. The targeted residues within the FP’s barrel were deduced based on comparison between the crystal structures of eGFP (1EMA), mKate (PDB 3bxb; for FusionRed; ^46^) and mScarlet (PDB 5lk4 ^39^) (Figure 4A and Suppl Table 2). Site directed libraries targeting seven homologous residues were created for FusionRed (A143, S144, N195, D197, R199, E201, and R221) and for mScarlet (A146, S147, N195, D197, K199, D201, and R221). Overall, for each template we screened a total of 322 mutants in four independent screens (1288 wells). The results of the functional testing of libraries from two templates showed similar trend. The majority of the screened variants (~930) showed complete lack of voltage sensitivity mainly due to loss of the fluorescence. However, several constructs based on both templates have shown doubling in signal amplitude (~4% %∆F/F vs. original ~2%) (Figure 4B).

**Figure 4.**
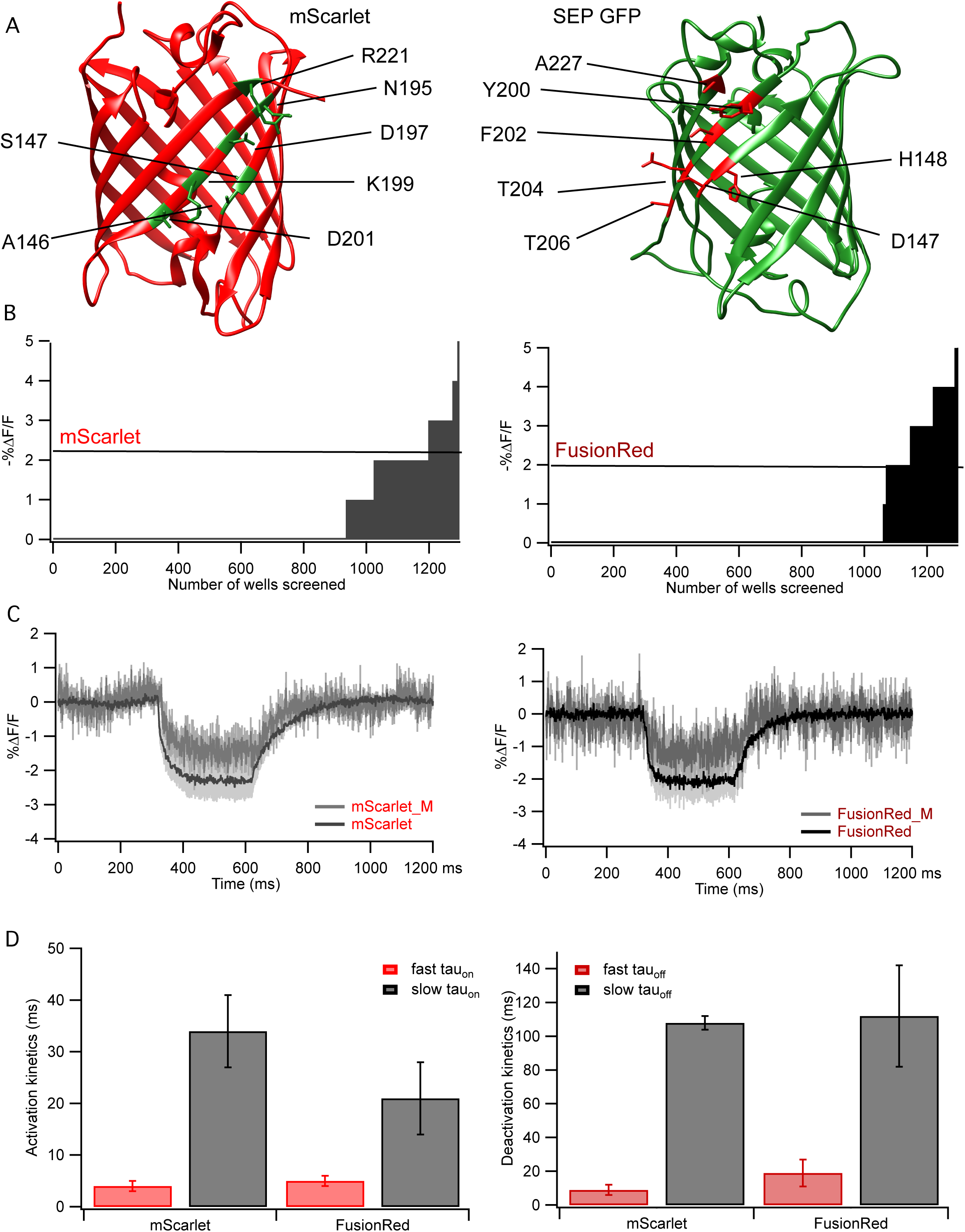
Modification of the beta barrel in the mScarlet and FusionRed GEVIs **A.** The directed mutagenesis of red-shifted FPs, mScarlet and FusionRed, was targeting residues know to affect the polarity and voltage sensitivity of GFP-based GEVIs. The crystal structures of eGFP (PDB:1EMA), mScarlet (PDB:5LK4) and mKate (PDB: 3bxb, not shown here) were used to identify structurally analogue residues to be mutated. The 3D structure of eGFP (PDB:1EMA) is used to show position of residues within SuperEcliptic pHluorin GFP (SEP GFP) that are relevant for voltage sensitivity of GEVIs ArcLight and Marina. **B.** We created site-directed mutagenic libraries targeted to seven residues (for details see Methods) in CiVSD-mScarlet and CiVSD-FusionRed GEVIs. For each library, 46 colonies were selected and tested in four independent screens for expression and voltage sensitivity. For each FP, the total of 1288 wells were screened. The functional testing of the libraries was performed on the custom-made, semiautomated screening platform which allows for simultaneous field stimulation and optical imaging in the 96-well dish format. The experiments were performed 24-48h postransfection using electrically active HEK293 cells transiently expressing various mutants. Each library was screened at last four times. The highest value for the voltage step-dependent fluorescence change in screens for parent construct (CiVSD-mScarlet and CiVSD-FusionRed) is indicated with the bar on the graph. **C.** The introduction of ArcLight-like mutations (analogs to ArcLight residues D147, F202, T204 and D227) in the red-shifted GEVIs has diminishing effect on voltage sensitivity. The combination of whole-cell patch clamp and optical imaging recordings were used for detailed characterization of CiVSD-mScarlet and CiVSD-FusionRed GEVIs. The experiments were done using non-electrically active HEK293 cells transiently expressing constructs. Each trace is average of 10 trials from four to six cells. All traces are unfiltered and corrected to remove photobleach effect. The SEM are shown as light colored traces. In left panel: CiVSD-mScarlet (dark grey) and CiVSD-mScarlet_M (FP mutant, light grey). In right panel: CiVSD-FusionRed (black) and CiVSD-mScarlet_M (FP mutant, dark grey). **D.** Kinetics of voltage-dependent response in red-shifted GEVIs, mScarlet and FusionRed. All data are shown as mean ± SEM. For each construct four to six cells were recorded. Left panel: activation speed. Right panel: deactivation speed.

In contrast, the combinatorial effect of mutations of the four residue homologues to the ones that attribute to the high sensitivity of ArcLight (D147, F202, T204 and D227) in both mScarlet and FusionRed GEVIs, tested with patch clamp electrophysiology, showed negative impact on voltage sensitivity causing for signal to decrease to ~-1%∆F/F compared to the GEVI(s) based on native FP(s) (Figure 4C and D).

## Material and methods

### Fluorescent protein selection

We tested a spectrally diverse pallet of FPs spanning over visible spectrum (excitation maximum 433-690 nm) derived from several different organisms. The blue-shifted variants are derivatives of either avGFP (*Aquorea victoria* GFP-mCerulean, eGFP, ecliptic pHluorin and eYFP) or lancelet *Branchiostoma lanceolatum* (mNeonGreen). For red-shifted proteins, we used either derivatives of *Discosoma striata* dsRED (mOrange, mOrange2, tdTomato, mTangerine, mStrawberry, and mCherry) or *Entacmaea quadricolor* eqFP580 (TagRFP, TagRFP-T, FusionRed, mNeptune, mNeptune2, and mCardinal). In addition, we tested recently developed synthetic FP mScarlet and near-red iRFP713, derived from acteriophytochrome RpBphP2 isolated from bacterium *Rhodopseudomonas palustris*.

#### GEVI design and construction

The initial constructs were created by replacement of fluorescent proteins (FPs) in the previously published *Ciona intestinalis* voltage sensitive phosphatase (CiVSP) based GEVIs, i.e. VSFP2.1 (insertion site R254 ^1^), Mermaid (insertion site S249 ^43^) and ArcLight (insertion site Q239 ^18^) with the range of single FPs. In all constructs CiVS contains point mutation R217Q that shifts its voltage sensitivity to a range more relevant for detection of neuronal electrical transients. The subcloning of FPs into respective vector(s) was done either by restriction enzyme (RE) digestion or by using In-Fusion PCR system for seamless cloning (Clontech, USA). Expression in HEK293 and mouse primary neuronal cells was facilitated by using vectors with either CMV or hSyn promoter, respectively. For RE digestion cloning, the FPs were PCR amplified using the *pfu* DNA polymerase (Stratagene, USA) with the coding sequences of the RE incorporated into the PCR primers. In the CMV vectors, the restriction enzymes used were either *NotI/EcoRI* for insertion at the fusion site S254 (as in VSFP 2.1), *BamHI/XbaI* for fusion sites Q239 (as in ArcLight; Addgene # 36856) and R249 (as in Mermaid). In some cases (mScarlet, FusionRed, and mNeonGreen) *EcoRI/SphI* sites were used for fusion at the site Q239 (as in Marina; Addgene # 74216). The enzyme digestion of the amplified DNAs was followed by the ligation reaction with the vector using T4 DNA ligase (BioNeer, USA). For seamless cloning of GEVIs into hSyn vectors, PCR amplification of the vectors and the FPs was performed using CloneAmp™ DNA polymerase (Clontech, USA). The PCR products were gel purified (Macherey-Nagel, USA) and used for ligation reaction. The ligation reaction was facilitated by introduction of 15bp overlapping extensions, identical to vector sequence at the insertion site, at each end of the FP during PCR and was performed using In-Fusion HD enzyme premix (Clontech, USA). In all cases, the DNA from ligation reactions was used to transform bacteria, colonies were picked and grown, and the DNA was extracted using miniprep kit (Macherey-Nagel, USA).

The point mutations and modifications in the linker sequence between CiVSD and FPs were done using the QuickCjhange II XL site-directed mutagenesis kit (Agilent Technologies, USA). The sequence verification of the novel constructs was done using Tag/dye-termination method (W. M. Keck Foundation, USA).

#### Library design and construction

The details of point mutation libraries production and functional testing in 96-well plate format has been previously described ^19^. In short, to facilitate functional screening of site-directed mutagenesis libraries CMV vectors for ArcLight Q239, mScarlet and FusionRed GEVIs were modified by GEVI fusion with a self-cleaving T2A peptide sequence (GSGEGRGSLLTCGDVEENPGP) followed by nuclear localized tag-FPs (mCherry in ArcLight; mCerulean in mScarlet and FusionRed). For production of the site-directed mutagenic libraries we used partially overlapping mutagenic primers during PCR vector amplification. For each PCR reaction, the mix of the three forward mutagenic primers containing one of the three degenerative codons (NDT, VHG or TGG) and the single reverse primer were used. The use of this combination of degenerative primers resulted in libraries encoding for all 20 amino acids with repeats for two amino acids Lys and Val. To facilitate subsequent vector circularization, the sequence for forward primers also included 15bp overlapping extensions with sequence identical to the vector sequence at the insertion site. For the PCR reaction we used CloneAmp™ DNA polymerase, followed by removal of the parent template by *DpnI* digestion. The ligation of amplified mutagenic vectors was done using In-Fusion HD enzyme premix (Clontech) and followed by bacterial transformation. For each site directed mutagenesis library, 46 bacterial colonies were selected (two libraries per 96-well plate; with four control wells) and cDNA was purified using automate liquid handling robot (epMotion 5057; Eppendorf, USA). The library complexity was confirmed by sequencing 10% of selected colonies.

#### Cell culture

This study was carried out in strict accordance with the recommendations in the Guide for the Care and Use of Laboratory Animals of the National Institutes of Health. The protocol was approved by the Pierce Animal Care and Use Committee.

The HEK293 cells (CRL-1573, ATCC, USA) were kept in the Dulbecco’s Modified Eagle Medium (High Glucose)-DMEM (Invitrogen, USA) supplemented with 10% fetal bovine serum-FBS (Sigma-Aldrich, USA). The electrically active HEK293 cells (CRL-3269, ATCC, USA) (Park et al., 2013) were kept in DMEM:F12 Medium (Invitorgen) supplemented with the 10% FBS, 0.5mg/ml G-418, and 2 ug/ml puromycin (Sigma-Aldrich). For the primary neuronal cultures, the hippocampal neurons were isolated from E18 mouse embryos and maintained in the culture medium containing Neurobasal medium, 0.5mM Glutamax-I, and B-27 supplement (Invitrogen). For the experimental assays, the HEK293 and primary neuronal cells were plated on the 12mm #1 coverslips coated with poly-D-lysine (Sigma-Aldrich). For the 96-well plate screening assays, the electrically active HEK293 cells were plated on the glass-bottom (#1) 96-well plates coated with poly-D-lysine. In all cases, cells were kept in the humidified incubator under control environmental conditions (at 37°C in atmosphere of 5% CO2). In all cases, for transient expression of GEVI constructs we used 0.1 µg per 96-well or 0.4 µg per 12 mm coverslip of DNA, and 0.25 µL per 96-well or 1 µL per per 12 mm coverslip of Lipofectamine2000 (Invitrogen). The experiments with the both lines of HEK293 cells were performed 24-48h post transfection. The primary neuronal cultures were used in experiments between DIV 10-14 (at least seven days post transfection).

#### Electrophysiology and fluorescence imaging

The recordings of membrane transients were performed in a perfused chamber with the bath temperature kept at 33-35°C by a temperature controller (Warner Instruments, USA). The bath solution was consistent of 150mM NaCl, 4mM KCl, 2mM CaCl2, 1mM MgCl2, 5mM D-glucose, and 5mM HEPES pH 7.4. We used a 3-5 MΩ glass patch pipettes (capillary tubing with 1.5/0.75 mm OD/ID) that were pulled on a P-97 Flaming/Brown type micropipette puller (Sutter Instrument Company, USA). The pipette solution contained 120mM K-aspartate, 4mM NaCl, 4mM MgCl2, 1mM CaCl2, 10mM EGTA, 3mM Na2ATP and 5mM HEPES pH 7.2. Voltage-clamp recordings in the whole-cell configuration were performed using either a Patch Clamp PC-505B amplifier (Warner Instruments, USA) or MultiClamp 700B amplifier (Molecular Devices,USA). The voltage stimulation protocols were written in LabView (National Instruments, USA). The voltage-dependent fluorescence signals were recorded in response to randomly applied depolarization or hyperpolarization voltage steps (20mV incremental steps, up to 100mV, 15 steps sweeps) starting at the holding potential of −70mV.

For recording of the hippocampal neurons, action potentials were initiated by 2ms constant current injection in whole-cell patch clamp mode. The pipette solution for neuron recordings contained (in mM): 120 K-gluconate, 3 KCl, 7 NaCl, 4 Mg2+-ATP, 0.3 Na-GTP, 20 HEPES and 14 Tris-phosphocreatin, pH adjusted with KOH to pH 7.3.

The cells were imaged using either Nikon Eclipse TE300 inverted microscope with a 60x 1.40 N.A. oil immersion lens or Olympus 51WI upright microscope with a 60x 1.10 N.A. water immersion objective. In most of the experiments, as a light source we used either an 150W Xenon arc lamp (OptiQuip, USA) or a SOLA Light Engine LED light source (Lumencor, Inc., USA). We used variety of the optical filter sets depending on the spectral characteristics of tested chromophores. Briefly, for mCerulean we used CFP-2432A filter set with a 438/24nm excitation filter, 458nm dichroic and 483/32nm emission filter. For GFP-based GEVIs we used 3035B filter set with a 472/30 nm excitation filter, a 495 nm dichroic mirror and a 520/35 nm emission filter. For eYFP and mNeonGreen we used YFP-2427A filter set (ex. 500/24nm, dic. 520nm, em. 542/27nm; Semrock Inc., USA). For the red-shifted FPs, we used following filter sets: Cy3-4040B (ex. 531/40nm, dic. 562nm, em. 593/40nm; Semrock), 49008 (ex. 560/40nm, dic. 585nm, em. 630/75nm), and 49006 cube (ex. 640/60nm, dic. 660nm, em. 700/75nm; Chroma Technologies, USA). The full list of optical filters used in this study can be found in the Suppl Table 1. For some of the experiments we used laser light, in which case for GFP chromophore we used a 488nm 50 mW laser (DL488-050, CrystaLaser, USA) and for RFPs chromophores 532nm 100mW laser (CL532-100). The laser light was transmitted to the microscope by a multi-mode fiber coupler (Siskiyou, USA), a quartz light guide and an Achromatic EPI-Fluorescence Condenser (Till Photonics, USA). Excitation light measured at the preparation was 8-18 mW/mm^2^, and was adjusted for each recording using a continuous circular neutral density filter (ThorLabs, Inc., USA) to the minimum required to record optical signals.

The fluoresce image was demagnified by an Optem® zoom system A45731 (Qioptiq LINOS, USA) and projected onto the 80×80 pixel chip of a NeuroCCD-SM camera controlled by NeuroPlex software (RedShirtImaging, USA). The images were recorded at a 1000-2000Hz frame rate. The measured traces were the spatial average intensity of all the pixels receiving light from the patched cell.

#### Libraries screening

The functional screening of the site-directed mutagenic libraries was done on a custom built semiautomated screening platform^19^. The platform was built around a Nikon Eclipse Ti-E inverted microscope equipped with a Perfect Focus System and a Nikon Plan Apo 20x 0.75 NA objective (Nikon, Japan). The light source used was either a SOLA Light Engine LED light source (Lumencor, Inc., USA) or pE-300 white CoolLED LED illumination System (CoolLED, Ltd., U.K.). A custom built imaging chamber attached to a motorized Prior Proscan II stage (Prior Scientific, Inc., USA) allows for constant temperature (37 °C) and humidity maintenance. For imaging we used Hamamatsu ORCA Flash 4.0 sCMOS camera (Hamamatsu, Japan) at a frame rate of either 50 or 100 Hz. The control plasmid used in all screenings was GFP-based GEVI ArcLight (Jin et al.; 2012). The filter set used for GFP GEVI ArcLight was a 472/30 nm excitation filter, 495 nm dichroic mirror and 520/35 nm emission filter (Semrock). For RFPs (nuclear localized mCherry in ArcLight, mScarlet and FusionRed in red-shifted GEVIs) we used a 560/40 nm excitation filter, 585 nm dichroic mirror and 630/75 nm emission filter (Chroma Technologies). For nuclear localized mCerulean in RFP-based GEVIs we used 438/24 nm excitation filter, 458 nm dichroic mirror and 483/32 nm emission filter (Semrock). The field stimulation of each well was facilitated by a custom made field electrode and actuator system attached to the roof of the imaging chamber. Electrical stimulation, imaging workflow and signal detection was controlled with a custom application written in LabView (National Instruments).

For cell detection, 16 images of the well bottom were collected and analyzed by a cell identification algorithm that quantified the fluorescence intensity of nuclear localized mCherry or mCerulean. The four fields with the largest number of cells were then selected by the program and used for analysis. For signal detection, a series of images were collected at 100 fps from the preselected fields of views using an appropriate filter set. During collection of each image series, a fixed pattern of electrical stimulation was applied to the cells. This routine was repeated until time-series images have been recorded from all 96 wells. The region-of-interest (ROI) mask for each cell within a field of view is created by a thresholding algorithm applied to the mCherry or mCerulean channel. A “cell mask” was generated by enlarging each of the FP nuclear features. A mean intensity trace for each cell is created from the time-series images recorded using the appropriate filter set. The magnitude of the fluorescence signal in each cell was quantified (ΔF/F), and various values of construct performance were evaluated to rank the signal size of a particular construct. Individual cells expressing the same construct showed large variation in fluorescence response amplitudes. We found it necessary to screen each construct in at least four separate plates to get an accurate picture of its performance. We also found that the maximum response amplitude across cells and wells was the most reliable indicator of particular constructs performance in subsequent patch clamp experiments. The values for resting light intensity of HEK cells expressing either GEVI were derived from mean intensity traces recorded using the appropriate filter set at the time points prior voltage stimulation. For each cell, resting light intensity value is average of three frames.

#### Confocal imaging

Confocal images of GEVI expressing cells were obtained with an Olympus Fluoview FV1000 (Olympus, USA) confocal laser scanning microscope equipped with LUMFL 60x/1.10 W objective. Depending on the imaged chromophore we used several lasers and filter sets. The 440nm diode laser and the BA480-495nm emission filter was used for mCerulean. A multi line Ar laser (458nm, 488nm, 515nm) was used for CFP in VSFP2.1(458nm line), and for GFP chromophores (eGFP, ecliptic pHluorin-488nm line). The emission filters used were the BA480-495nm and the BA505-525nm, respectively. For the red-shifted chromophore (mOrange, mKok, mStrawberry, mCherry and TagRFP) we used 543nm HeNe(G) laser and BA560-660nm emission filter. For image acquisition and processing FV10-ASV confocal software was used.

#### Data analysis

The image sequences, current, and voltage traces were acquired using NeuroPlex software (RedShirt Imaging). The further data analysis was done using custom programs written in LabView (National Instruments) and Igor (Wavemetrics, USA). Images of all cell types were recorded at a frame rate of 500, 1000 or 2000 Hz, and depicted optical traces were simple spatial averages of intensity of all pixels within the region of interest (ROI). The fluorescence traces were corrected for the photobleaching by dividing the signal by a single or double exponential curve fitted to the portion of the trace that does not contain a voltage step.

Data are presented as percent change in ∆F/F.

The probe dynamics were fitted with either a single exponential equation,

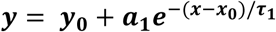

or a double exponential equation,

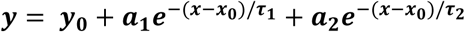

The ΔF/F versus V plot was analyzed with the Boltzmann equation:

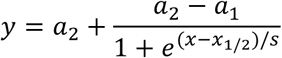

where *a*_1_ and *a*_2_ are asymptotic values for % fluorescence change at small and large x (membrane potential, mV), respectively. τ_1_ and τ_2_ are time constants in ms, x_1/2_ is membrane potential at half maximum ΔF/F and S is steepness of the curve (mV).

## Discussion

Protein evolution based on highthroughput mutagenesis and functional screening significantly improved our ability to develop and optimize GEVIs ^11, 13, 19, 47^. Such efforts would be significantly enhanced by an understanding of how the primary protein sequence contributes to the physical rearrangements that modulates voltage-dependent fluorescence output.

One popular voltage sensitive fluorescent indicator design is replacement of the downstream enzyme domain of voltage sensitive phosphatase ^41^ with a fluorescent protein. This substitution is downstream of the S1-4 domain. In these indicators the voltage induced fluorescent signal stems from conformational changes in fluorescent protein(s) caused by movements of the S1-4 domains. We and others ^13, 18, 19, 21, 23–25, 44, 48, 49^ have extensively studied this design in efforts to produce indicators with maximum sensitivity, differing spectral properties and increased response speed. We have constructed and tested a wide range of VSP-fluorescent protein constructs and examined their responses to voltage steps ^18, 19, 21, 23, 44^. To our surprise these experiments, most exemplified by the current study, reveal a paradox related to VSD GEVI performance. We have found that although discrete changes (point mutations) in the fluorescent protein have had enormous effects on the voltage sensitivity of indicators (i.e. GEVI Arclight and Marina; ^18, 19^), using fluorescent proteins with little primary sequence homology, from highly divergent organism, with different chromophores, biochemical characteristic and spectral properties produce nearly identical indicators that show depolarization-dependent decrease in fluorescence ^48^.

The activity dependent change of fluorescence intensity in FP-biosensors can result from changes in the extinction coefficient, quantum yield, and/or change in the protonation equilibrium of the chromophore ^50^(Molina et al., 2019). From existing studies, it seems that amino acid residues surrounding the chromophore have the most effect on amplitude of GEVI voltage sensitivity ^13, 18, 19, 23, 24^. Changes in VSD sequence are more likely to influence the kinetics of voltage response ^24, 26, 44, 51^. However, both sensitivity and kinetics have been improved by modification of linker sequence that connects the VSD and FP ^18, 44^. While various environmental factors can influence FP output, the fluorescence change in VSD-bound FPs is not likely to be the consequence of cytosolic pH fluctuations caused by neuronal transients ^48, 52, 53^.

Previously we showed that high sensitivity of the GEVI ArcLight can be attributed to outward oriented charged residues (D147, F202, T204 and D227) that populate surface of *β* barrel (beta sheets 7, 10 and 11) of SuperEcliptic pHluorin GFP ^52^. In addition, we showed that further mutagenesis of ArcLight (residues D147A, H148A and Y200V) yields reverse in the signal polarity without loss of dynamic range (GEVI Marina ^19^). However, the exact mechanism behind voltage sensitivity of ArcLight and Marina is still not clear. The existing hypothesis contribute voltage induced spatial re-arrangement of the FP residues to change in hydrophobic interactions at the FP surface upon translocation relative to the plasma membrane ^19^, intramolecular dimerization ^52, 53^, and/or mechanical distortion of FP ^54^. Here, we showed that targeted mutagenesis of homologous residues in red-shifted FPs (mScarlet and FusionRed) does not produce significant changes in voltage sensitivity. Unlike with the GFP-based GEVIs, we showed that combinatorial effect of mutations of these residues lowers the voltage sensitivity compared to GEVI based on native FPs. In a recent study, the potential of dimerization of the FP for GEVI development was explored using red-shifted dimer FP dTomato ^45^.

However, the resulting GEVI Ilom shows similar performance as GEVIs based on tandem dimer tdTomato described here. In addition, similar to our results introduction of charged outward oriented residues on the FP surface failed to increase the dynamic range of Ilom sensitivity ^45^.

Currently there are no illustrative model(s) of the structure/function interactions that explain how VSD-based fluorescence voltage indicators work. The optimal approach for model development would be to examine all structural components of GEVIs (VSD, FP and linker in-between) on molecular level while systematically manipulating transmembrane potential, from resting to various activated states (depolarization and hyperpolarization). Unfortunately, considering membrane localization of the voltage transients this is experimentally very challenging, if not impossible. The biophysical studies allow for simultaneous recording of gating currents and fluorescent signals ^26, 43^. This approach offers insight into correlation between VSD conformational dynamics and fluorescence output, but not on the atomic level dynamics of GEVI components. Consequently, the most valuable insights on GEVI performance and function are still gained via empirical studies.

To this end, our study in which we systematically examined the most promising red-shifted FPs for their potential to detect voltage transients contributes to our overall understanding of VSD GEVI function. These FPs vary by fluorescence spectra which is the result of the structural differences in the chromophore and/or the coordinating amino acids. Surprisingly our results show that major alterations in primary sequence do not affect the basic characteristics of the relationship between membrane voltage and fluorescent light output. This signifies that the biophysical change that occurs with membrane potential deflections has the same effect on light output regardless of the chromophore or primary amino acid sequence of the FP. The use of various FPs in the CiVS-FP design mostly results in GEVIs with voltage response characteristics that vary in a narrow range, insufficient to report the naturally occurring variety of voltage transients in neurons. This insight coupled with the fact that single amino acid changes that produce radical improvements in sensitivity (i.e. Arclight and Marina) strongly supports the empirical approach of finding functional constructs and then performing directed protein evolution coupled with functional screening to drive the desired change in the response properties.

## Acknowledgements

The authors are grateful to Lei Jin for providing preliminary data for the study, and Marko A. Popovic for technical assistance with imaging experiments. We are grateful to Malcolm Connor and Chelsea Gardiner for the help in performing library screening experiments. We are grateful to Larry Cohen for helpful discussions and advice. We would also like to thank The John B. Pierce Laboratory, Inc. for ongoing support. We would like to acknowledge the expert technical contributions of the Pierce Laboratory Instrument shop including John Buckley, Andrew Wilkins, Tom D’ Alessandro, Ronald Goodman and Angelo DiRubba. The authors have no competing interests to report.

This study was supported by NIH grants U24 NS057631, U01 NS090565, R01 NS083875 and U01NS103517.

**Suppl_Table 1.** Optical and molecular properties of fluorescent proteins that were used for creation of GEVIs described in this study.

**Suppl_Table 2.** Amino acid residues targeted in directed evolution of red-shifted GEVIs. Amino acid residues numbering in SuperEcliptic pHluorin (SEP GFP) in GEVI ArcLight, and GEVI Marina, mScarlet and FusionRed.

## Notes

### Competing Interest Statement

The authors have declared no competing interest.

