## Supplementary material for "Different categories of fluorescent proteins result in GEVIs with similar characteristics": Suppl_Tables

| Parent FP | Fluorescent protein | Excitation (nm) | Emission (nm) | ε[c] ($10^3$ $M^{-1}cm^{-1}$) | QY | pKa | Oligomerization | Filter set used in this study | Reference |
|---|---|---|---|---|---|---|---|---|---|
| avGFP | mCerulean | 433 | 475 | 43 | 0.62 | 4.7 | Monomer | **Exc.:** 438/24<br>**Dic.:** 458<br>**Em.:** 483/32 | Rizzo et al., 2004 |
| | EGFP | 488 | 507 | 56 | 0.60 | 6.0 | Weak dimer | **Exc.:** 472/30<br>**Dic.:** 495<br>**Em.:** 520/35 | Nagai et al., 2002 |
| | Ecliptic pHluorin | * | * | * | * | 7.1 | Monomer | | Miesenbock et al., 1998 |
| | EYFP | 513 | 527 | 67 | 0.67 | 6.9 | Weak dimer | **Exc.**: 500/24<br>**Dic.**: 520<br>**Em.**:542/27 | Nagai et al., 2002 |
| LanYFP | mNeonGreen | 506 | 517 | 116 | 0.8 | 5.7 | Monomer | | Shaner et al., 2013 |
| dsRED | mOrange | 548 | 562 | 71 | 0.69 | 6.5 | Monomer | **Exc.:** 531/40<br>534/42<br>560/40<br>**Dic.:**560<br>562<br>585<br>**Em.:** 593/40<br>605/75<br>630/75 | Shaner et al., 2004 |
| | mOrange2 | 549 | 565 | 58 | 0.60 | 6.5 | Monomer | | Shaner et al., 2004 |
| | tdTomato | 554 | 581 | 138 | 0.69 | 4.7 | Pseudo-monomer | | Shaner et al., 2004 |
| | mTangerine | 568 | 585 | 38 | 0.3 | 5.7 | Monomer | | Shaner et al., 2004 |
| | mStrawberry | 574 | 596 | 90 | 0.29 | <4.5 | Monomer | | Shaner et al., 2004 |
| | mCherry | 587 | 610 | 72 | 0.22 | <4.5 | Monomer | | Shaner et al., 2004 |
| eqFP578 | TagRFP | 555 | 584 | 100 | 0.48 | <4.0 | Monomer | | Merzlyak et al., 2007 |
| | TagRFP-T | 555 | 584 | 81 | 0.41 | 4.6 | Monomer | | Shaner et al., 2008 |
| | FusionRed | 580 | 608 | 94.5 | 0.19 | 4.6 | Monomer | | Shemiakina et al., 2012 |
| | mNeptune | 600 | 651 | 75 | 0.23 | 5.4 | Monomer | **Exc.:** 605/55<br>**Dic.:** 660<br>**Em.:** 680/30<br>700/75 | Lin et al., 2008 |
| | mNeptune2 | 599 | 651 | 89 | 0.24 | 6.3 | Monomer | | Chu et al., 2014 |
| | mCardinal | 604 | 659 | 87 | 0.19 | 5.3 | Monomer | | Chu et al., 2014 |
| N/A | mScarlet | 564 | 594 | 100 | 0.70 | 5.3 | Monomer | **Exc.:** 560/40<br>**Dic.:** 585<br>**Em.:** 630/75 | Bindels et al., 2016 |
| rpBphP2 | iRFP (iRFP713) | 690 | 713 | 105 | 0.059 | 4.0 | Dimer | **Exc.:** 640/60<br>**Dic.:** 660<br>**Em.:** 700/75 | Filonov et al., 2011 |

| SEP GFP (ArcLight) | SEP GFP (Marina) | mScarlet | FusionRed |
|---|---|---|---|
| D147 | A147 | A146 | A143 |
| H148 | A148 | S147 | S144 |
| Y200 | V200 | N195 | N195 |
| F202 | F202 | D197 | D197 |
| T204 | T204 | K199 | R199 |
| T206 | T206 | D201 | E201 |
| A227 | A227 | R221 | R221 |
